# Sonogenetic stimulation of the brain at a spatiotemporal resolution suitable for vision restoration

**DOI:** 10.1101/2021.11.07.467597

**Authors:** S. Cadoni, C. Demené, M. Provansal, D. Nguyen, D. Nelidova, G. Labernede, I. Alcala, J. Lubetzki, R. Goulet, E. Burban, J. Dégardin, M. Simonutti, G. Gauvain, F. Arcizet, O. Marre, D. Dalkara, B. Roska, J. A. Sahel, M. Tanter, S. Picaud

**Affiliations:** Sorbonne Université, INSERM, CNRS, Institut de la Vision, 17 rue Moreau, F-75012 Paris, France; Physics for Medicine Paris, INSERM, CNRS, École Supérieure de Physique et de Chimie Industrielles (ESPCI Paris), Paris Sciences et Lettes (PSL) Research University, 75012 Paris, France; Institute of Molecular and Clinical Ophthalmology Basel, Basel, Switzerland; Department of Ophthalmology, The University of Pittsburgh School of Medicine, Pittsburgh, PA 15213, United States; Department of Ophthalmology and Vitreo-Retinal Diseases, Fondation Ophtalmologique Rothschild, F-75019 Paris, France; Centre Hospitalier National d’Ophtalmologie des XV-XX, F-75012 Paris

**Author notes:** These authors contributed equally to this work (M. Tanter, S. Picaud).

## Abstract

Remote, precisely controlled activation of the brain is a fundamental challenge in the development of brain-machine interfaces providing feasible rehabilitation strategies for neurological disorders. Low-frequency ultrasound stimulation can be used to modulate neuronal activity deep in the brain^1–7^, but this approach lacks spatial resolution and cellular selectivity and loads the brain with high levels of acoustic energy. The combination of the expression of ultrasound-sensitive proteins with ultrasound stimulation (‘sonogenetic stimulation’) can provide cellular selectivity and higher sensitivity, but such strategies have been subject to severe limitations in terms of spatiotemporal resolution *in vivo*^8–10^, precluding their use for real-life applications. We used the expression of large-conductance mechanosensitive ion channels (MscL) with high-frequency ultrasonic stimulation for a duration of milliseconds to activate neurons selectively at a relatively high spatiotemporal resolution in the rat retina *ex vivo* and the primary visual cortex of rodents *in vivo*. This spatiotemporal resolution was achieved at low energy levels associated with negligible tissue heating and far below those leading to complications in ultrasound neuromodulation^6,11^. We showed, in an associative learning test, that sonogenetic stimulation of the visual cortex generated light perception. Our findings demonstrate that sonogenetic stimulation is compatible with millisecond pattern presentation for visual restoration at the cortical level. They represent a step towards the precise transfer of information over large distances to the cortical and subcortical regions of the brain via an approach less invasive than that associated with current brain-machine interfaces and with a wide range of applications in neurological disorders.

---

It was anticipated that brain-machine interfaces (BMIs) based on multi-electrode arrays would provide solutions for many neurological disorders or deficits, including blindness^12^. Multi-electrode arrays have met with increasing success in peripheral sensory system rehabilitation strategies, for restoring hearing in the cochlea or sight in the retina, for example. The restoration of vision is, perhaps the most demanding challenge for BMIs, as it ultimately requires the video-rate transmission of complex spatial patterns of stimulation. In patients who have lost the eye-to-brain connection, great hopes have been raised for the restoration of sight by cortical multi-electrode arrays since early reports in the 1960s^13,14^.However, a recent clinical study with surface cortical electrodes showed that form perception required sequential point-by-point stimulation for more than one second for letter recognition^15^. The cortical surface electrodes used (0.5 mm in diameter) had a wide spacing (2-4 mm), limiting pixel numbers^15^ and precluding their use for complex tasks, such as navigation and face recognition. Studies in non-human primates have shown that stimulation deep within the visual cortex is much more efficient for eliciting perception with smaller currents^16^. With the use of penetrating electrodes, non-human primates were able to recognize letters following the static multi-point electrical stimulation of their visual cortex for only 167 ms^17^. The higher temporal resolution associated with this cortical vision strategy was achieved at the expense of invasiveness, due to the need for penetrating electrodes. Clinical trials with this technology are underway, but a loss of efficacy over time has already been reported^14^. Nevertheless, these results demonstrate the feasibility of visual restoration at the cortical level through stimulation deep within the cortex.

Optogenetic therapy provides an alternative for the non-invasive stimulation of neurons at higher resolution, as demonstrated on the retina^18–21^. However, despite encouraging preliminary results in studies aiming to elicit visual perception at the cortical level^22–24^, approaches for optical stimulation of the cortex are hindered by the dura mater and by the scattering and absorption of light by tissues. Penetrating light guides have therefore been proposed as a means of maintaining the resolution of optical stimulation and its intensity within the brain after removal of the dura^23–25^. Ultrasound (US) waves could potentially overcome these limitations, making it possible to focus at depth and to achieve the non-invasive neuromodulation of cortical and subcortical areas of the brains ^1–7^. The implantation of ultrasonic matrix arrays in a cranial window would provide a less invasive approach to vision restoration, as the ultrasonic beam would propagate through the intact dura, the subdural and subarachnoid spaces (Fig. 1a). Unfortunately, existing US neuromodulation strategies are restricted to low-frequency transmissions, resulting in poor spatial resolution (>3 mm) and long-lasting responses. Indeed, attempts at high-frequency neuromodulation have resulted in high levels of acoustic energy^26^, with a risk of thermal heating^8-10^ and US-mediated tissue damage^6,11^. This inevitable trade-off between spatial resolution and acoustic intensity has greatly limited the applicability of US neuromodulation for BMI. The sonogenetic therapy approach proposed here aims to solve this problem. The ultimate goal is the development of a technology that is less invasive than electrodes but capable of activating neurons in the visual cortex with a high spatial (∼100 µm) and temporal (<50 ms) resolution. We propose 1) to boost neuronal sensitivity to US through the expression of US-sensitive channels on cell membranes^8–10,27,28^, 2) to demonstrate that it is possible to target a locally defined subset of neurons by gene therapy, which is not currently possible with non-specific US neuromodulation strategies, 3) to induce responses with a sufficiently high temporal precision and 4) to gain more than one order of magnitude in spatial resolution through the use of high-frequency US, which was previously considered impossible *in vivo* without considerably increasing acoustic intensities and potential adverse effects^29^. This activation of the brain with a unique combination of high spatial and temporal resolution, through the intact dura, could render sonogenetic therapy perfectly compatible with applications for vision restoration, which require video-rate patterns of stimulation.

**Fig. 1.**
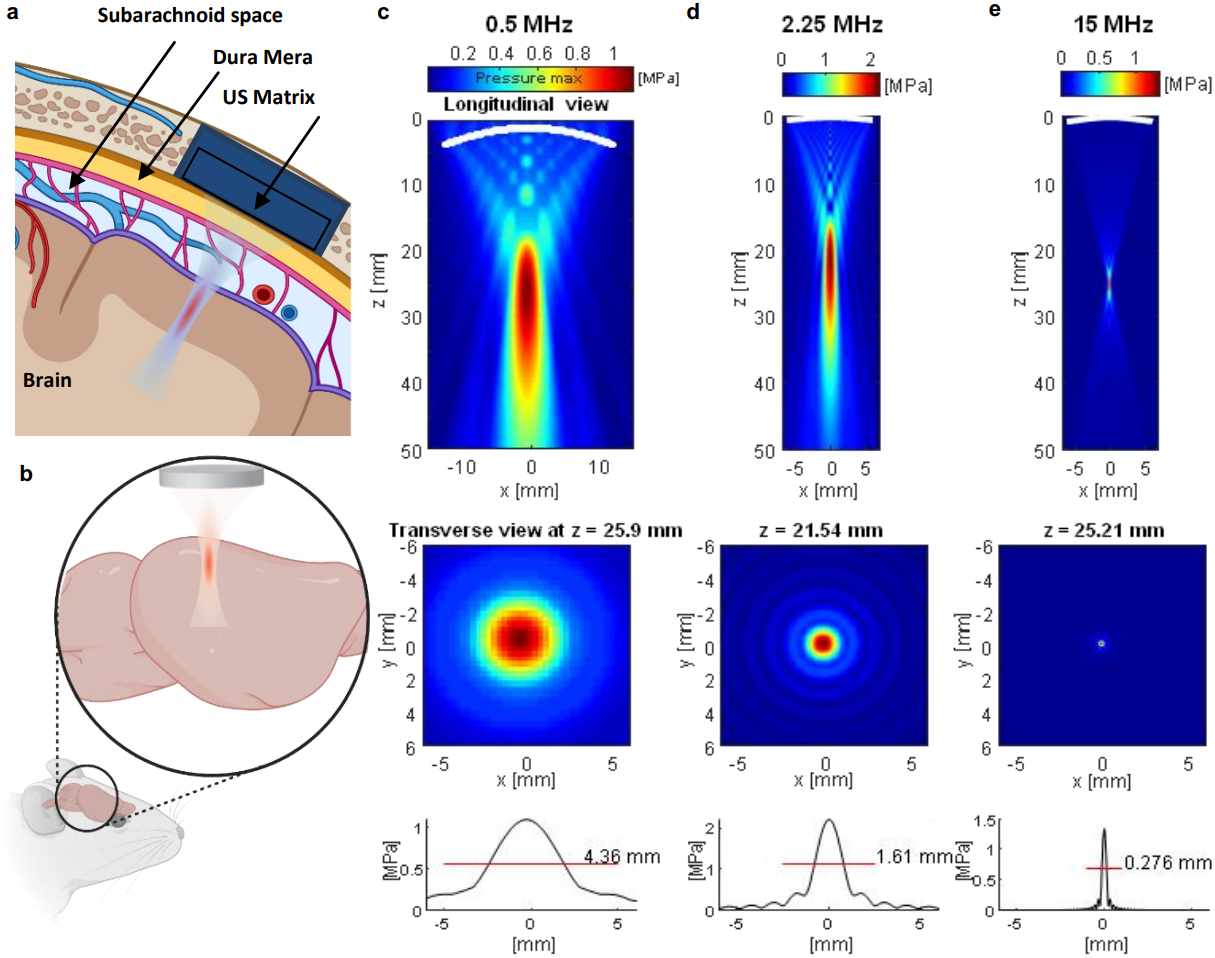
Sonogenetics using focused ultrasound beams for visual restoration through the intact dura mater: impact of ultrasonic transmision frequency. (a) Concept of visual restoration with US matrix arrays implanted in a cranial window for localized US neuromodulation of the primary visual cortex in humans. The US beam can be adaptively focused at different locations in the V1 cortex while passing through the intact dura mater, subdural and subarachnoid spaces. (b) Proof-of-concept setup used in this study for V1 sonogenetic activation in rodents, using a high-frequency focused transducer on a craniotomized mouse. (c) Characterization of the radiated field for the 0.5 MHz transducer used in this study. (top) Longitudinal view of the maximal pressure for a monochromatic acoustic field radiated at 0.5 MHz by the 25.4 mm Ø, 31.75 mm focus transducer. Pressure maximum is reached at 25.9 mm, slightly closer to the transducer than the geometric focal point, which is a documented effect ^66^. (middle) Transverse section of the maximal pressure field at depth z = 25.9 mm. (bottom) One-dimensional profile of this transverse section giving the FWHM of the focal spot (4.36 mm at 0.5 MHz). (d) Same characterization for the 2.25 MHz 12.7mm Ø 25.4 mm focus transducer. (e) Same characterization for the 15 MHz 12.7mm Ø 25.4 mm focus transducer. Note that the maximum pressure is reached very close to the geometric focus (25.21 mm versus 25.4 mm for the geometric focus) for this configuration. The FWHM of the focal spot is 0.276 mm.

With the aim of developing a suitable approach to demonstrate proof-of-concept for sonogenetic brain activation, we first characterized three focused ultrasonic transducers. Their dimensions and geometric foci were selected so as to provide a relevant model of future implanted matrix arrays for human applications and to be suitable for use in proof-of-concept experiments in rodents (Fig. 1b). Transducers were designed with a similar focal distance (F = 31.7 mm for the lower frequency and F = 25.4 mm for the two higher frequencies) and numerical apertures (diameter Ø = 25.4mm for the lower frequency and Ø = 12.7 mm for the 2 higher frequencies, corresponding respectively to an aperture number F/Ø= 1,25 and 2 respectively), for the transmission of focused beams over different frequency ranges (*f = 0*.*5 MHz, f = 2*.*25 MHz* and *f = 15 MHz*, corresponding to wavelengths of 3.0, 0.7 and 0.1 mm, respectively) (Fig. 1c-e). Increasing the frequency of ultrasound stimulation from 0.5 MHz (typical of neuromodulation) (Fig. 1c) to 15 MHz (Fig. 1e) has a major impact on the resolution that can be achieved: in our case the volume of the focal spot and therefore of the stimulated locus is diminished by a factor ∼4100 when replacing the 0.5 MHz transducer by the 15 MHz transducer. We therefore performed most of our experiments at 15 MHz; the two lower frequencies were used for an initial comparison of efficiency and spatial resolution.

Before investigating cortical activation *in vivo*, we studied sonogenetic therapy in the retina, which serves as an easily accessible part of the central nervous system. In this natural mammalian neuronal circuit, we expressed the mechanosensitive ion channel of large conductance (MscL)^30–34^ specifically in rat retinal ganglion cells (RGCs), with *in vivo* intravitreal delivery by an adeno-associated vector (AAV). Here, RGCs were simply used as a neuronal model suitable for the *ex vivo* screening of acoustic parameters for effective sonogenetic stimulation in a realistic neural circuit, avoiding potential US interference with the auditory system *in vivo*^35,36^. Vectors were produced with the MscL gene from *Escherichia coli* in its wild-type (WT) form and with a mutation, G22S^37^, which increased the sensitivity of cultured neurons to mechanical and ultrasonic stimulation^34,38^. An AAV2.7m8^39^ serotype vector was used to encode the MscL channels fused to the red fluorescent protein tdTomato, under control of the SNCG promoter to target the RGC population specifically among the retinal neurons^40^. Following AAV injections, tdTomato expression was detected *in vivo*, on the eye fundus (Fig. 2a). Examination of the flat-mounted retina showed that tdTomato expression was restricted to the ganglion cell layer and the optic fiber bundles (Fig. E1b). We further demonstrated that expression was limited to RGCs, by labeling these cells with a specific antibody, RPBMS (Fig. 2b). Expression of the MscL gene seemed to be concentrated at the cell membrane on the soma and axon (Fig. 2c). The staining indicated that, in the transfected area, 33.73% and 45.83% of RPBMS-positive cells expressed tdTomato, for the MscL-WT and MscL-G22S proteins, respectively (Fig. 2d).

**Fig. 2.**
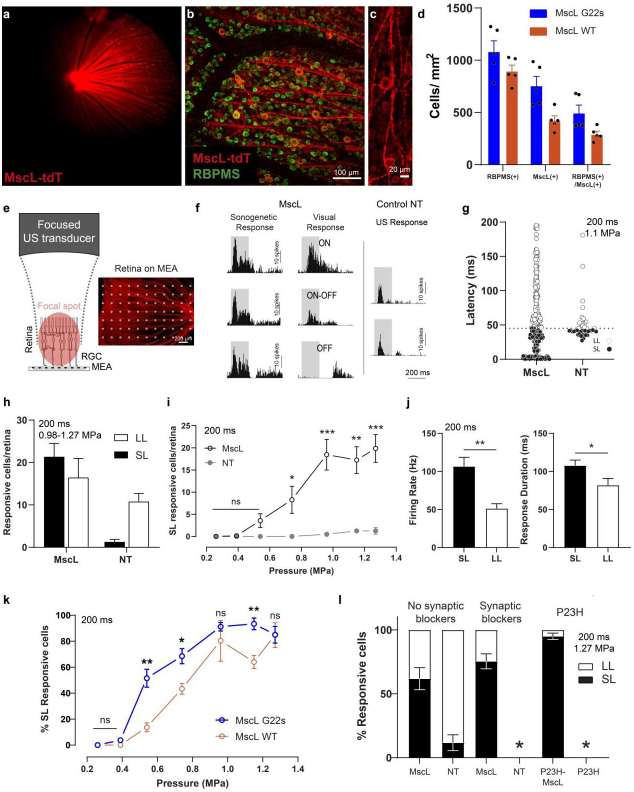
Sonogenetic therapy in retinal ganglion cells. (a) Retinal fundus image showing MscL-tdTomato expression in the *in vivo* rat retina. (b) Representative confocal stack projection across the RGC layer of a flat-mounted retina expressing MscL-tdTomato (red) and labeled with anti-RBPMS antibody (green). (c) Magnification of a few RGCs expressing MscL-tdTomato, showing its expression at the soma and axonal levels of the membrane. (d) Density of RBPMS-positive, MscL-positive and double-labeled cells for 5 MscL G22S and 5 MscL WT retinas. (e) (Left) Schematic diagram of the experimental setup for focused US stimulation of the isolated retina on a multielectrode array (MEA). (Right) Retina expressing MscL-tdTomato on a MEA chip. White dots represent electrodes. (f) Representative peristimulus time histograms (PSTH) for RGCs, for different stimuli. (Left) PSTH of three RGCs expressing MscL, showing a short latency and a sustained response after the start of a 15 MHz US stimulus (1.27 MPa). (Middle) Corresponding PSTH of the same RGCs in response to a visual stimulus, showing typical ON, ON-OFF and OFF responses. (Right) PSTH of two RGCs from a non-transfected (NT) retina in response to a 15 MHz US stimulus (1.27 MPa), showing a response to the start and end of the stimulus. (g) RGC latencies in response to a 15 MHz US stimulus for MscL (*n*=300 cells, 9 retinas) and NT (*n*=41 cells, 4 retinas) retinas. The dotted line represents the 45 ms threshold that divides cells into short latencies (SL: black dots) and long latencies (LL: white dots) cells. (h) Mean number of cells per retina responding to 15 MHz US stimuli (0.98-1.27 MPa) for MscL (*n*=9) and NT (*n*=4) retinas and for LL and SL cells. (i) Mean number of SL-responding RGCs per retina following stimulation with US stimuli of increasing pressure for MscL (*n*=9) and NT (*n*=4) retinas.*, *p*=.0356, **, *p*=.0010, ***, *p*=.0008, unpaired *t* test. (j) Mean maximum firing rate and mean response duration of SL and LL RGCs from MscL retinas in response to 15 MHz stimuli of increasing pressure (0.2-1.27 MPa) (n=9, **, *p*=.0017, *, *p*=.0418, unpaired *t* test). (k) Percentage of SL RGC cells (normalized against the maximum number of responsive cells in the experiment) responding to US stimuli of increasing pressure for MscL WT (*n*=3) and MscL G22S (*n*=6) retinas. *, *p*=.0173, **, *p*=.0065, **, *p*=.0083, unpaired *t* test. (l) Percentage of RGCs responding to US stimulation, for retinas in normal conditions (*n*=9 retinas for MscL and 4 for NT), and following the application of a cocktail of synaptic blockers (CNQX-CPP-LAP4) (*n*=3 retinas each for MscL and NT), and for P23H retinas with (*n*=3 retinas) and without (*n*=3 retinas) MscL expression. The ratio of SL to LL is shown for each set of conditions.

We then evaluated RGC sensitivity to US, by performing *ex vivo* recordings of the retina on a multi-electrode array (Fig. 2e). In retinas expressing the MscL channel, RGCs displayed strong and sustained ON responses to focused 15 MHz US stimulation (Fig. 2f-left). These responses were different from that recorded in non-transfected (NT) retinas, in which RGCs presented an increase in spiking activity after the start of the stimulus (Fig. 2f-right) with relatively long latencies, 50.4 ± 4.2 ms (Fig. 2g). By contrast, many RGCs of transfected retinas presented responses with a very short latency, 12.2±2.5 ms, (Fig. 2f-left), whereas others continued to respond with longer latencies (Fig. 2g). RGCs were classified into SL and LL in terms of their response, with SL corresponding to a latency of less than 45 ms. Even if some RGCs of MscL-expressing retinas continued to respond with longer latencies, they presented a significantly shorter overall geometric mean latency, 24.35 ms (GSD: 6.06) than non-transfected RGCs, 46.62 ms (GSD: 1.42) (*p*<.0001, unpaired *t*-test on log-transformed values) (Fig E2c-d). The generation of SL ON US responses was not related to a specific RGC type (Fig. E2a), as they were measured both in cells with ON responses to light, and those with OFF responses to light (Fig. 2f-left). MscL expression decreased latency and increased the mean number of cells per retina responding to US (Fig. 2h). SL responding cells expressing MscL were sensitive at much lower US pressures than non-transfected cells and their number increased with increasing US pressures (Fig. 2i). SL US responses also involved higher firing rates and were more sustained than LL US responses (Fig. 2j). Moreover, we observed that the G22S mutation further enhanced the sensitivity of SL RGCs to lower US pressures (Fig. 2k).

We further investigated the origin of the sonogenetic responses, by adding a mixture of synaptic blockers (CNQX-LAP4-CPP) to the bath perfusing the retina. These synaptic blockers were found to abolish US responses in non-transfected retinas but not in MscL-transfected retinas, in which they did not suppress the SL US responses and only decreased the number of LL US responses (Fig. 2l, Fig E2c-d). This observation suggests that the MscL-mediated SL US responses are initiated by MscL-transfected RGCs, whereas LL US responses in the transfected and non-transfected retina originate upstream from RGCs. These conclusions were supported by data recorded with the retinas of blind P23H rats. No US responses were recorded for the non-transfected P23H retina (Fig. 2l), demonstrating that the LL US responses required synaptic transmission, as previously reported^41^, and suggesting a possible origin in photoreceptors. MscL-transfected P23H retinas displayed many SL US responses and few LL US responses, further demonstrating that MscL expression generates the SL responses in RGCs (Fig. 2l, Fig E2c-d). The residual LL responses for P23H and for blocker-bathed retinas may originate from RGCs with a lower level of MscL expression or from US wave reverberations in the recording chamber. The geometric mean latencies in all the MscL-tested groups were already very different from those for the non-transfected retina (Fig. E2c), but the cumulative distribution of latencies further highlighted these differences between the non-transfected retina and the other tested conditions in terms of latency distribution (Fig. E2d). We subsequently restricted our analyses to SL US responses. We investigated the temporal kinetics of US responses under various durations of US stimulation (Fig. 3a) and at various repetition rates (Fig. 3b). Neurons responded to even very short stimulation durations (10 ms), with responses showing a fast return to the control level of activity (Fig. 3a). For longer stimuli (100 ms or longer), habituation began to occur, with a reduction of the maximum firing rate (Fig. 3c). US response durations were correlated with stimulus duration (Fig. 3d). Using a different repetition rate of a 15 MHz US stimulus, RGCs were able to follow the rhythm up to a stimulus repetition frequency of 10 Hz (Fig. 3b-e). The Fano factor in the previous experiments indicated that the response had a low variability in spike count and possibly high information content (Fig. 3c-e).

**Fig. 3.**
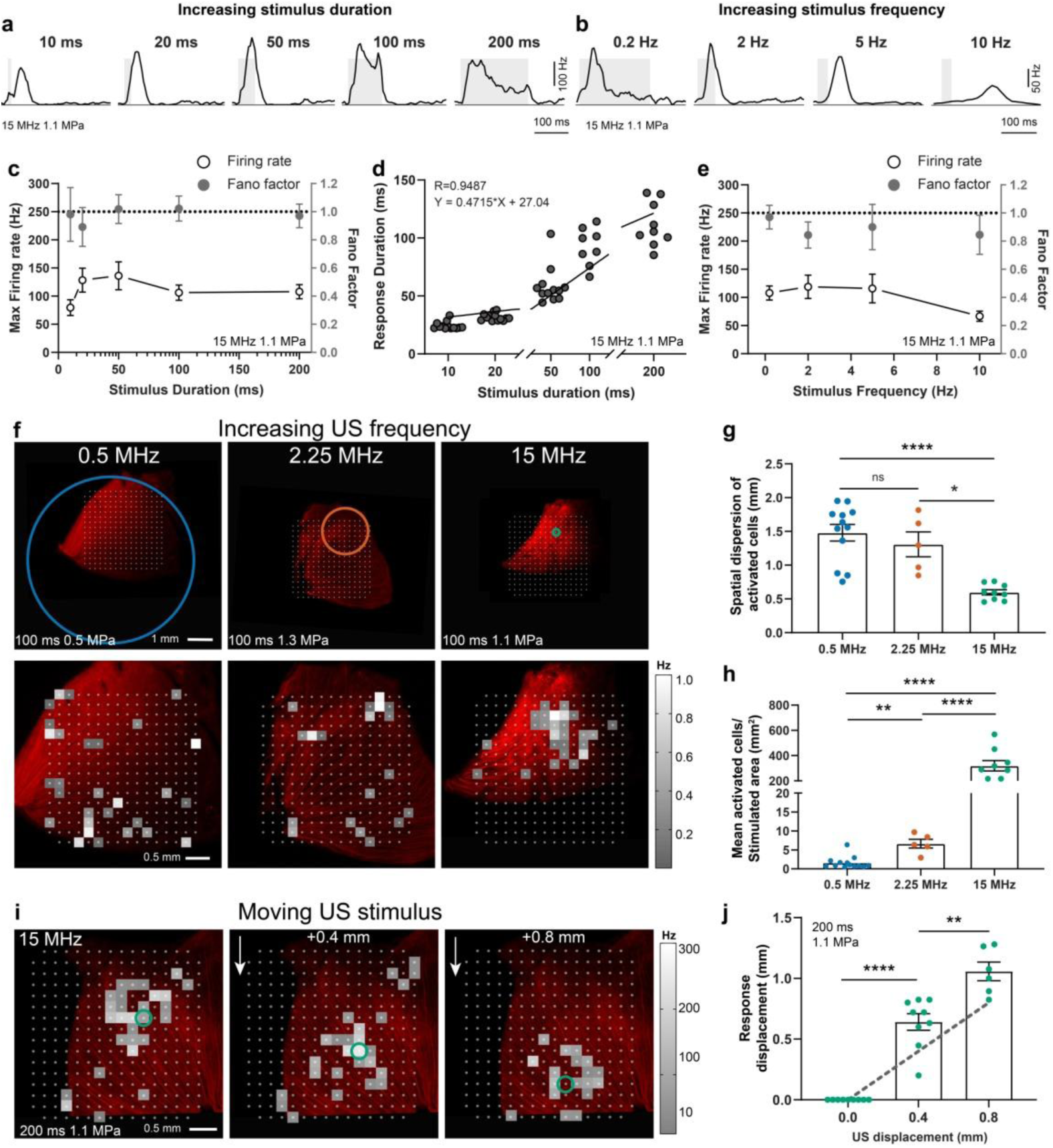
Spatiotemporal properties of sonogenetic retinal responses. (a-b) Spike density functions of two representative RGCs from a *MscL*-expressing retina for various 15 MHz stimulus durations (0.5 Hz stimulus repetition rate) (a) and stimulus repetition frequencies (stimulus durations: 10, 20, 50, 200 ms) (b).(c) Mean maximum firing rate for different 15 MHz stimulus durations and mean Fano factor values for all cells (*n*=9 retinas, except for stimulus durations of 10 and 20 ms *n*=8).(d) Correlation between response duration and stimulus duration, confirmed by the linear regression line (*n*=9 retinas). (e) Mean maximum firing rate for different stimulus repetition frequencies and mean Fano factor values for all cells (*n*=9 retinas, except for stimulus frequencies of 5 and 10 Hz *n*=8). (f) (Top) Retinas on a MEA chip and corresponding size of the incident US pressure beam (circles represent the FWHM and are centered on the estimated center of response), for 0.5, 2.25 and 15 MHz. (Bottom) Corresponding activation maps following US stimulation. Each square box represents an electrode with at least one US-activated cell. Color map representing the normalized firing rates of the cells. (g) Spatial dispersion of activated cells calculated as the mean Euclidean distance between activated cells weighted according to maximum firing rate and (h) ratio of the number of activated cells to the area stimulated on the MEA chip for the three US frequencies. ****, *p*<.0001, **, *p*=.0008,*, *p*=.0169, unpaired *t* test. *n*=12 retinas for 0.5 MHz (0.29-0.68 MPa), *n*=5 retinas for 2.25 MHz (1.11-1.62 MPa), *n*=9 retinas for 15 MHz (1.12-1.27 MPa). (i) Heatmaps showing the activated cells of a *MscL-*transfected retina following a relative displacement (0, +0.4 and +0.8 mm) of the 15 MHz US transducer. Each colored box represents an activated cell; the color map shows the maximum firing rate. Circles represent the estimated center of the response. (j) Relative displacement of the center of the response following displacement of the 15 MHz US transducer. ****, *p*<.0001, **, *p*=.0018, unpaired *t* test. *n*=9, 9, and 6 positions for 4, 4 and 2 retinas for displacements of 0, 0.4 and 0.8 mm, respectively. The dotted gray line represents the theoretical displacement.

We then investigated whether different US frequencies (0.5, 2.25 and 15 MHz) affected the spatial resolution of the response, in accordance with the measured US pressure fields, which became smaller at higher US frequencies (Fig. 1b-d, Fig. E3). The features of the responses evoked by the different US frequencies were found to be similar (Fig. E2e-f). Figure 3f illustrates the distribution of responding cells on the recording chip under different US stimulation frequencies with the expected FWHM (full width at half maximum) distribution of the US pressure field (colored rings). Cells responding to US were widespread over the recorded area for 0.5 and 2.25 MHz, but appeared to be more confined for 15 MHz (Fig. 3f). For each stimulated retina, we then calculated the spatial dispersion of activated cells; this value decreased significantly from 1.48±0.12 mm and 1.30±0.18 mm at 0.5 MHz and 2.25 MHz, respectively, to 0.59±0.03 mm at 15 MHz (Fig. 3g). These distances were consistent with the size of the measured ultrasound pressure fields (Fig. 1c-e); for the 0.5 MHz transducer, the focal spot of which was much larger than the MEA chip. This distance is close to the mean distance between randomly selected pairs of electrodes on the MEA chip (1.73 mm). The density of activated cells increased significantly with increasing US frequency and activated cells were more widely dispersed on the larger stimulated area at lower frequencies (Fig. 3h). US stimulation is more effective at higher frequencies, because lower acoustic power values are required to activate an equivalent number of cells. Indeed, even if the acoustic intensities at 2.25 and 15 MHz were quite similar, the acoustic power delivered was almost two orders of magnitude lower at 15 MHz (0.03 W) than at 2.25 MHz (0.82 W). Interestingly, at 15 MHz, the stimulated area was small enough for the focal spot of the US probe to be moved above the isolated retina, triggering a shift in the responding cells (Fig. 3i). This shift followed the probe’s focal spot over the retina, in the same direction and with a consistent displacement amplitude. The center of the response was found to move in accordance with the displacement of the US transducer (Fig. 3j). These results demonstrate that our sonogenetic therapy approach can efficiently activate neurons with a millisecond and sub-millimetric precision.

Following this *ex vivo* demonstration on the retina, we then investigated whether the approach could also be applied to the brain *in vivo*, paving the way for a sonogenetic BMI using high frequency ultrasonic arrays implanted in the skull bone. As the G22S mutation enhanced the US sensitivity of RGCs *ex vivo*, we expressed this channel in the cortical neurons of the primary visual cortex (V1) in rats. We injected a AAV9.7m8 vector encoding the MscL-G22S channel fused to tdTomato under the control of the neuron-specific CamKII promoter into V1. TdTomato fluorescence was detected in the brain (Fig. 4a) and in cortical slices (Fig. 4b). V1 neurons expressed tdTomato, particularly in layer 4 (Fig. 4b). Staining with an anti-NeuN antibody showed that 33.4% of cortical neurons in the transfected area expressed tdTomato (Fig. 4c).

**Fig. 4.**
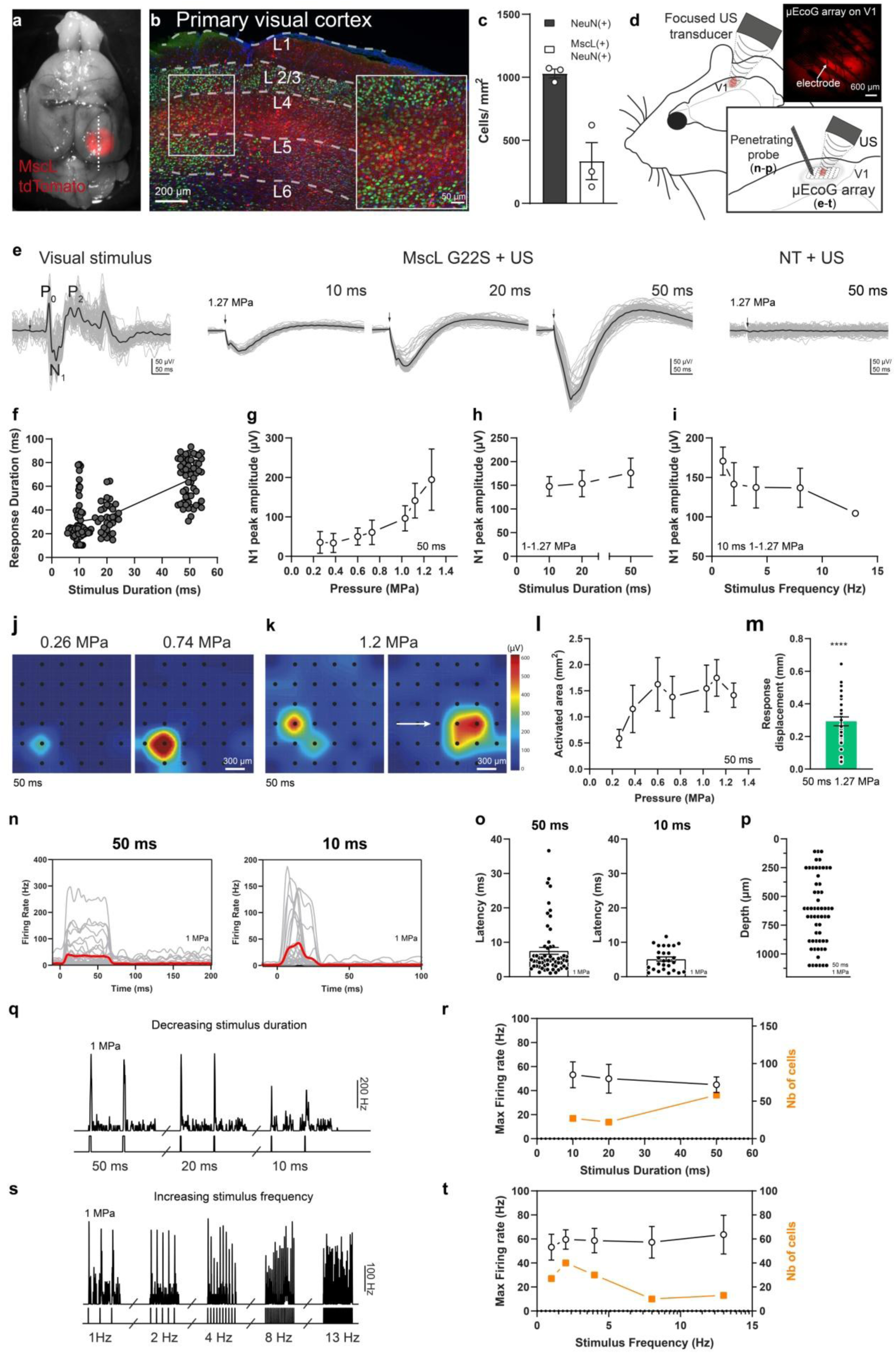
*In vivo* sonogenetic therapy in V1 cortical neurons. (a) Image of a rat brain expressing MscL-G22S-tdTomato (red) in V1. The dotted line illustrates the location of a sagittal slice. (b) Representative confocal stack projection of a sagittal brain slice expressing MscL G22s-tdTomato (red) and labeled with anti-NeuN antibody (green) and DAPI (blue). The layers of V1 are delineated by dashed white lines. (Lower right) Magnification of layer 4 of V1. (c) Density of NeuN-positive, MscL-positive and double-labeled cells for 3 brain slices. (d) Schematic diagram of the setup used for *in vivo* electrophysiological recordings and US stimulation; (Top right) µEcoG electrode array placed on V1 of a *MscL*-transfected rat. (e) (Left) Representative visual-evoked cortical potentials in response to a 100 ms flash. (Middle) Representative sonogenetic evoked potentials for 15 MHz US stimuli of various durations. (Right) Representative responses of a non-transfected (NT) rat to a 15 MHz US stimulus. Black traces represent the mean evoked potential over 100 trials. Each gray trace represents one trial. The black arrow indicates the start of the stimulus. (f) Duration of sonogenetic µEcog responses for stimuli of different durations (10 ms *n*=58, 20 ms n=32 and, 50 ms n=56 trials on 6 animals). (g) N1 peak amplitude for increasing US pressure, (i) increasing duration and (j) frequency (*n*=6 rats). (J) Pseudocolor activation maps for stimuli of increasing US pressure and (k) for a horizontal displacement of the US transducer by 0.8 mm (the arrow indicates the direction of the displacement). Each black dot represents an electrode of the electrode array. The color bar represents N1 peak amplitude in µV. (l) Area activated for various US pressure values (*n*=6 animals). (m) Displacement of the activation center relative to the previous position following movement of the US transducer by 0.4 mm. *p*<.0001, one-sample *t* test, *n*=37 positions on 6 animals. (n) Spike density functions (SDF) of 58 and 27 neurons following US stimulation for 50 and 10 ms, respectively, for MscL rats. (o) Response latencies following 50 and 10 ms US stimuli (50 ms *n*=58 cells, 7 rats; 10 ms, n=27 cells, 5 rats). (p) Depth of US-responding cells (*n*=58) in *MscL*-expressing rats (*n*=7). (q) Instantaneous SDF of responses to US stimuli of different durations (1 Hz stimulus repetition frequency). (r) Maximum firing rates and numbers of activated neurons upon US stimulations of different durations (US pressure: 1 MPa). (s) Instantaneous SDF of responses to US stimuli of different repetition frequencies (10 ms stimulus duration). (t) Maximum firing rate and number of activated neurons upon US stimulation at different stimulus repetition frequencies (10 ms, 1MPa).

For investigation of the ability of a 15 MHz US stimulus to activate cortical neurons, we placed a micro-EcoG (µEcoG) electrode array on the cortical surface of V1 (Fig. 4d). In non-transfected (NT) animals, no US-evoked signal was recorded (Fig. 4e-right, *n*=3 rats), whereas, in V1 expressing MscL-G22S, US stimulation of the cortical surface elicited large negative µEcoG potentials (Fig. 4e-middle, *n*=6 rats). These US-evoked negative deflections were different from the recorded visual-evoked potentials, which presented typical P0, N1 and P1 positive and negative deflections (Fig. 4e-left). The duration of the US responses was clearly related to the duration of the US stimulation (Fig. 4f). The amplitude of US-evoked potentials increased with both increasing US pressure (Fig. 4g) and increasing US stimulus duration (Fig. 4h). V1 cortical responses were again able to follow a repetition rate of up to 13 Hz (Fig. 4i) even if peak amplitude decreased slightly for increasing stimulation frequencies.

We then investigated the spatial distribution of US-evoked neural activity. The peak depolarization of each channel was measured and linearly interpolated to build pseudocolor activation maps (Fig. 4j). The size of the US-responding cortical area was dependent on the US pressure (Fig. 4j-k), and ranged from 0.58 ± 0.17 mm^2^ (*n*=6 rats) to 1.41 ± 0.23 mm^2^ (*n*=5 rats) for US pressures of 0.26 and 1.27 MPa, respectively (Fig. 4l). We investigated the possibility of achieving patterned US stimulations, by moving the US transducer in 0.4 mm steps over the recorded area. When the ultrasound probe was moved laterally, the source of the generated neuronal activity moved in a similar direction (Fig. 4k). The spatial location of the evoked potentials moved, on average, by 0.29 ± 0.09 mm (*n*=6 rats) from the previous location (Fig. 4m), even though we moved the US transducer in 0.4 mm steps. These measurements were probably conditioned by the 300 µm discrete spatial pitch distribution of the electrodes and the lateral spread of activity in the circuit. These results suggest that our approach to sonogenetic therapy could yield a spatial resolution of within 400 µm for stimulations at 15 MHz, the focal spot of our 15 MHz transducer being 276 µm wide (Fig. 1d). This opens up the possibility of targeting small areas (down to 0.58 mm^2^ for 0.26 MPa), depending on the pressure level. These very localized US-evoked responses and their dependence on the position of the US probe confirmed that they were due to the activation of MscL-G22S-expressing neurons and not to an indirect response related to auditory activation, as previously suggested by others^35,42^.

The possibility of using US stimulation to activate neurons at different depths was then explored. V1 neurons were recorded with a 16-site penetrating multi-electrode array (Fig. 4d). In V1 neurons expressing MscL-G22S, US stimulation at 15 MHz generated sustained responses even to 10 ms-long US stimuli (Fig. 4n). The latency of these responses was short (5.10 ± 0.62 ms n=27 cells and 7.51 ± 1.00 ms n=58 cells, for 10 ms and 50 ms stimuli respectively, Fig. 4o), consistent with direct US activation of the recorded cortical neurons. Responding neurons were recorded at various cortical depths, ranging from 100 µm to 1 mm (Fig. 4p), the focal spot diameter of the US probe being 3.75 mm in the xz plane. Deep neurons responded reliably to stimuli of decreasing duration, from 50 ms to 10 ms, with similar firing rates to stimuli of different durations, whereas longer stimuli induced responses in a broader population of neurons (Fig. 4q-r). By increasing stimulus frequencies to up to 13 Hz, we were able to evoke responses in cortical neurons (Fig. 4s) with equivalent firing rates, but the number of responding cells decreased with increasing stimulus frequency (Fig. 4t).

We investigated whether sonogenetic stimulation could also induce light perception, by assessing mouse behavior following 15 MHz US stimulation of V1 in MscL G22S-transfected (*n*=9) (Fig. E5a) and non-transfected (*n*=7) animals. Mice subjected to water deprivation were trained to associate the visible-light stimulation of one eye with a water reward, as previously described ^43^ (Fig. 5a). This task was learned within four days, as indicated by the increasing success rate during this period, from 23.57% to 76.09% (Fig. 5b). The success rate was determined by assessing the occurrence of an anticipatory lick between the light being switched on and the release of the water reward 500 ms later (Fig. 5a). Following this associative learning phase, the mice were subjected to US stimulation of V1 on day 5 (<u>Fig.</u> b). Following US stimulation at the highest pressure, MscL-G22S-transfected mice achieved a success rate (66.98%) similar to that following light stimulation on day 4 (76.10%) (Fig. 5b). After a pause during the weekend (day 6-7), the animals had partially forgotten the task associating sonogenetic stimulation with a water reward (Fig. 5b). However, they rapidly recovered an efficient association on day 10 (Fig. 5b). We found that the latency of the first anticipatory lick was shorter for sonogenetic stimulation (193.2 ± 12.8 ms; *n*=9) than for stimulation with a light flash (285.3 ± 12.4 ms; *n*=15) (Fig. 5c). This shorter latency for the US response is consistent with the faster activation of cortical neurons for sonogenetic stimulation than for light stimulation of the eye (Fig. 4e), suggesting a shorter delay in the transfer of visual information from the eye to V1. Non-transfected animals were unable to associate the US-stimulation of their cortex with the water reward (Fig. 5d), demonstrating that the sonogenetic activation of cortical neurons was truly the triggering factor, as opposed to the auditory perception of US, as previously reported for low-frequency US stimulation^35,42^. We applied different US pressures to the visual cortex in transfected mice and found that success rate increased with pressure (Fig. 5d). Interestingly, the licking frequency during the 500 ms before delivery of the water reward also increased with US pressure (Fig. 5e). These results indicate that the sonogenetic stimulation of the visual cortex generates light perception in mice.

**Fig. 5.**
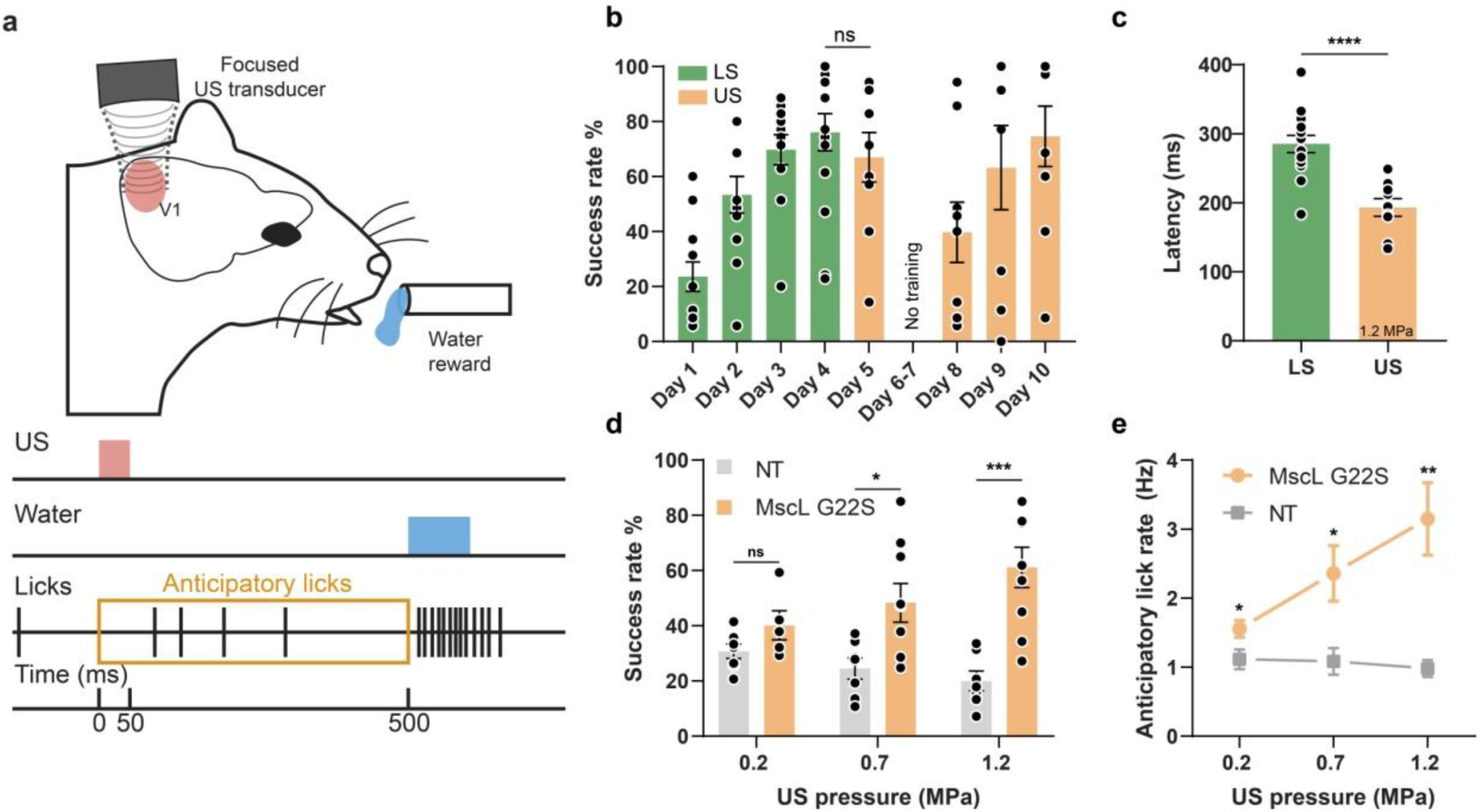
Behavioural response induced by sonogenetic activation of the V1 cortex in mice following associative visual training. (a) Schematic diagram of the behavioral task performed by mice. Water-restricted animals trained in an associative learning paradigm for light stimulation with a water reward are subjected to US stimulation of VA at 15 MHz. A trial is considered successful if the animal performs at least one anticipatory lick within the 500 ms time lag between stimulus onset and water reward. (b) Mean rates of successful trials for 4 days of training during learning of the association between light stimulation (50 ms) and water reward (LS green) followed by the US stimulation (US orange) (50 ms 1.2 MPa, ns, *p*=.4311, unpaired *t* test). (c) Time to first lick after light (50 ms) and US stimulation (50 ms, 1.2 MPa) (****, *p*<.0001, unpaired *t* test). (d) Mean rate of successful trials over 4 days of US stimulation for non-transfected (NT) and MscL-G22S transfected mice, following 50 ms of US stimulation at increasing US pressure (ns, *p*=.0751, *, *p*=.0114, ***, *p*=.0006, unpaired *t* test, for 0.2, 0.7 and 1.2 MPa, respectively). (e) Anticipatory lick rates for NT and MscL-G22S transfected mice at increasing US pressures (*, *p*=.0424, *, *p*=.0150, **, *p*=.0031, unpaired *t* test, for 0.2, 0.7 and 1.2 MPa, respectively).

The development of remotely controlled cortical and subcortical deep neuronal stimulation techniques is of considerable interest for the treatment of diverse neurological diseases and sensory handicaps. Optogenetics has been developed for this purpose in non-human primates, but its potential for transfer into clinical practice is limited by the low level of light penetration into the brain tissue^23,24^. US stimulation can overcome this limitation^2,44– 46^, but the optimal conditions for sonogenetic approaches, in terms of both genetic constructions and acoustic parameters, are far from clear. Most previous sonogenetic studies focused on the use of low-frequency US^8–10^ as, in particular, in the recent demonstration of MscL-based sonogenetic activation in mice brain with long bursts of low-frequency US^10^. However, such low-frequency US waves lead to limited centimetric spatial resolutions (typically around 5×5×45 mm^3^) and an uncontrolled spatial beam distribution, due to the formation of standing waves^47^. In the rodent brain, this phenomenon generates reverberations throughout the entire braincase^48^, with the probable activation of non-target structures, such as the auditory pathway^35,42^. An alternative approach to spatially containing the US stimulation involves the use of higher US frequencies, but this was thought to demand higher energy levels, exceeding safety limits and favoring tissue damage^26^. The bacterial MscL channel has been reported to sensitize neurons to US^10,38^ and to lower the pressure for neuronal activation, but its use for high-spatiotemporal resolution sonogenetic stimulation has yet to be shown to be effective *in vivo*. We first demonstrated the expression of MscL *in vivo* in the neurons of the retina. We then determined the most appropriate conditions for activating the MscL-transfected retinal neurons with high-frequency US with an appropriate temporal and spatial precision. In the absence of MscL expression, the retina can respond to US, but with different temporal characteristics, due to the intrinsic mechanosensitivity of photoreceptors^41^. This attribution of the natural US response to photoreceptors was supported by its absence in the retinas of P23H rats presenting severe photoreceptor degeneration or following synaptic blockade in non-transfected retina (Fig. 2l). Due to the reported rewiring in the P23H retina, we cannot totally exclude another origin for the natural response to US. However, previous studies showing electrical activation of the P23H rat retina via the retinal network are not consistent with this possibility^49–51^. Working towards our goal of cortical visual restoration, we then showed that MscL-G22S was effectively expressed in cortical neurons. The activation of these MscL-expressing neurons resulted in responses with millisecond latencies and a spatial resolution of at least 400 µm in the *xy* plane and 3.75 mm on the *z* axis, consistent with neuronal responses throughout the depth of the cortex (Fig. 4m-p). Moreover, MscL G22s expression in the visual cortex led to a behavioral response upon US stimulation, confirming light perception following sonogenetic activation of the visual cortex. These sonogenetic responses were genuinely related to MscL expression, as they were not observed in non-transfected animals. The sonogenetic approach presented here greatly decreased the US pressure required for the activation of RGCs and V1 cortical neurons by high-frequency US with stimulation sequences remaining below FDA safety limits (510k, Track 3) for US imaging (e.g. for a 10 ms US stimulus of 0.6 MPa, the Isptp is 12 W/cm^2^ and the Ispta value is 0.12 W/cm^2^). These very low acoustic pressures and acoustic intensities prevent tissue damage, as they are similar to those that have been widely used in clinical diagnostic imaging for decades^52,53^ Moreover, simulations of US-induced heating in brain tissue revealed that typical US parameters (i.e. 20 ms, 1.27 MPa) (Fig. 4e-h) increased the local temperature by an estimated 0.12 °C (see methods), with even high repetition rates (up to 13 Hz) leading to a moderate temperature increase (<0.3 °C) (Fig. E4c-f). These low-temperature fluctuations and stimulation sequences compliant with FDA limits suggest that our approach had no toxic side effects and that US-elicited responses were not temperature-driven and were therefore probably mediated by mechanical activation of the MscL channel by US. Following on from previous demonstrations that the MscL channel is a suitable sonogenetic actuator^10,34,38^, we provide *in vivo* evidence that the MscL channel has appropriate kinetics for the activation of neurons at a precise spatiotemporal resolution *in situ* and *in vivo*. Previous *in vivo* studies reported the restoration of form vision at cortical level with 0.5 to 1 mm surface electrodes spaced more than 1 mm apart^13–15^. The resolution of the proposed sonogenetic therapy therefore appears to be compatible with the restoration of form vision. Moreover, the resolution of the approach could be increased by using gene therapy to drive expression in specific cell populations and cell compartments^54^. Further studies are required to generate an interface for coding visual information into US patterns transmitted by an ultrasonic matrix array onto the visual cortex at a video rate. Computer algorithms will probably be required to model complex visual features, such as flicker fusion, in the US encoder. However, our approach provides great hope for the development of high-resolution visual restoration at the cortical level, through its unique combination of a rapid response, high spatial resolution, and some cell selectivity with promoters, all features essential for video-rate brain-wide pattern stimulation. Even if this approach requires craniotomy, as for other existing visual prostheses, it provides a less invasive approach based on deep and distant cortical activation from above the dura mater following AAV cortical injections. The reported long-term expression of genes following AAV gene therapy^55^ suggests that this strategy could provide stable sight restoration. More generally, it paves the way for a new type of brain-machine interface capable of compensating for disabilities and suitable for use in the treatment of neurological disorders.

## Methods

### Animals

All experiments were conducted in accordance with the National Institutes of Health Guide for the Care and Use of Laboratory Animals. The experimental protocols were approved by the Local Animal Ethics Committee (Committee Charles Darwin no. 5, registration number 9529 and 26889) and conducted in agreement with Directive 2010/63/EU of the European Parliament. All rats included in this study were Long Evans rats from Janvier Laboratories or P23H (line 1) transgenic rats. Wild-type mice (C57BL/6J) were obtained from Janvier Laboratories. Before being included in experimental procedures, at least 5 days of adaptation to the local animal facility were allowed to animals in which they were enclosed in a controlled environment with a half-day dark/light cycle with nutrition *ad libitum* with enrichment.

### Plasmid cloning & AAV production

Plasmids containing the *Escherichia coli MscL* sequence in the WT form and with the G22S mutation were obtained from Francesco Difato (Addgene plasmids #107454 and #107455)^56^. For the targeting of retinal ganglion cells, the SNCG promoter^40^ was inserted into an AAV backbone plasmid containing the *MscL* sequence fused to the tdTomato gene and the Kir2.1 ER export signal, to drive expression at the plasma membrane. An AAV2.7m8 vector was used for intra-vitreal delivery. For the targeting of neurons in the cortical layers of V1, the SNCG promoter was replaced by the CamKII promoter and an AAV9.7m8 vector was chosen. All the recombinant AAVs used in this study were produced by the plasmid cotransfection method, and the resulting lysates were purified by iodixanol purification as previously described, to yield high titer recombinant AAV virus ^57^.

### US stimulus

Three focused ultrasound transducers with different central frequencies were used, to obtain focal spots of different sizes: a low-frequency transducer at 0.5 MHz (diameter Ø = 1 inch = 25.4mm, focal distance f = 1.25 inch = 31.7 mm) (V301-SU, Olympus), a transducer at an intermediate frequency of 2.25 MHz (Ø = 0.5 inch = 12.7 mm, f = 1 inch = 25.4 mm) (V306-SU, Olympus) and a high-frequency transducer at 15 MHz (Ø = 0.5 inch = 12.7 mm, f = 1 inch = 25.4 mm) (V319-SU, Olympus). Acoustic fields radiated by those three focused transducers are presented in Figure 1 (simulations) and extended figure E3 (experimental measurements). A TiePie Handyscope (HS3, TiePie Engineering) was used to produce the stimulus waveform, which was then passed through an 80 dB RF power amplifier (VBA 230-80, Vectawave) connected to the transducer. Transducer pressure outputs (pressure at focus, 3D pressure maps) were measured in a degassed water tank with a Royer-Dieulesaint heterodyne interferometer^58^. The US stimuli used for *ex vivo* and *in vivo* stimulation had the following characteristics: 1 kHz pulse repetition frequency with a 50% duty cycle, sonication duration between 10 and 200 ms and inter-stimulus interval between 0.01 and 2 s. Peak acoustic pressures were ranging from 0.11-0.88 MPa, 0.3-1.6 MPa, 0.2-1.27 MPa, for the 0.5, 2.25 and 15 MHz transducers, respectively. The corresponding estimated Isppa values were 0.39-25.14 W/cm^2^, 2.92-83.12 W/cm^2^ and 1.30-52.37 W/cm^2^.

### Ex vivo

#### Intra-vitreal gene delivery and retinal imaging

Rats were anesthetized with isoflurane (5% for induction, 3% for maintenance) and 2 µl of AAV suspension, containing between 8 and 14 × 10^10^ viral particles, was injected into the center of the vitreous cavity with direct observation of the tip of the needle. One month after injection, fluorescence imaging was performed on the injected eyes, with a Micron IV retinal imaging microscope (Phoenix Research Laboratories) used to observe MscL expression via the fluorescent tdTomato tag. Electrophysiological recordings were performed at least one month after injection.

#### MEA recordings

Retinas were isolated under dim red light, in Ames’ medium (A1420, Sigma-Aldrich) bubbled with 95% O_2_ and 5% CO_2_ at room temperature. Pieces of the retina were flattened on a filter membrane (Whatman, GE Healthcare Life Sciences) and placed on a poly-L-lysine (0.1%, Sigma) coated multi-electrode array (electrode diameter 30 µm, spacing 200 µm, MEA256 200/30 iR-ITO, MultiChannel Systems) with retinal ganglion cells facing the electrodes. The retina was continuously perfused with bubbled Ames medium at 34 °C, at a rate of 2 ml/min, during experiments.

TdTomato fluorescence was checked before recordings, by using a stereo microscope (SMZ25, Nikon) to observe transgene expression in the recorded area. For some experiments the AMPA/kainate glutamate receptor antagonist 6-cyano-7-nitroquinoxaline-2,3-dione (CNQX, 25 μM, Sigma-Aldrich), the NMDA glutamate receptor antagonist [3H]3-(2-carboxypiperazin-4-yl) propyl-1-phosphonic acid (CPP, 10 μM, Sigma-Aldrich) and a selective group III metabotropic glutamate receptor agonist, L-(+)-2-amino-4-phosphonobutyric acid (L-AP4, 50 μM, Tocris Bioscience), were freshly diluted and bath-applied through the perfusion line 10 minutes before recording. Full-field light stimuli were delivered with a digital micro-mirror display (DMD, Vialux, resolution 1024×768) coupled to a white light LED light source (MNWHL4, Thorlabs) focused on the photoreceptor plane. An irradiance of 1 µW/cm^2^ was used. The US transducers were coupled with a custom-made coupling cone filled with degassed water, mounted on a motorized stage (PT3/M-Z8, Thorlabs) and placed orthogonally in the recording chamber above the retina. For positioning of the US transducer over the retina, the reflected signal of the MEA chip and the retina was detected with an US-key device (Lecoeur Electronique). The distance between the retina and the transducer was equal to the focal length of the transducer; this was verified with the flight time of the reflected signal. RGC recordings were digitized with a 252-channel preamplifier (MultiChannel Systems). Spikes from individual neurons were sorted with SpykingCircus software^59^. RGC responses were then analyzed with custom scripts written in Matlab (MathWorks). They were classified as ON, ON-OFF or OFF, with the response dominance index^60^. The latency of each cell relative to the start or end of the stimulus was calculated as the time between the start of the stimulus and the maximum of the derivative of spike density function. For cells responding to US stimulation, two classes were identified on the basis of latency — short and long latency — by fixing a threshold equal to the minimum of the latency distribution of the responses of non-transfected cells to US (45 ms). We determined the peak value A of spike density function for the calculation of response duration, which was defined as the time interval between the two time points for which the SDF was equal to A/*e* (*e*: Euleur’s number). The percentage of cells responding to US stimulation of increasing US pressure was calculated as the ratio of the number of activated cells to the maximum number of responding cells for all the US pressures considered. The Fano factor, quantifying spike-count variability, was calculated as the ratio of the variance of the spike-count to the mean. Values close to 1 indicate that information can be transmitted. The Euclidean distance between two activated cells was weighted according to the maximum firing rate of the cells. The ratio of the number of activated cells to the size of the area stimulated on the MEA chip was calculated considering the size of the US focal spot for 2.25 and 15 MHz and the size of the MEA for 0.5 MHz, because the focal spot was larger than the MEA for this frequency. The center of the response was estimated by weighting the maximum firing rate of each cell by its distance from other responding cells, and the displacement of the response was calculated as the Euclidean distance between two center-of-response positions.

### In vivo

#### Intracranial injections

Rats or mice were anesthetized with a ketamine/medetomidine mixture (40 mg/kg / 0.14 mg/kg) or ketamine/xylazine (80 mg/kg / 8 mg/kg) respectively, and injected subcutaneously with Buprenorphine (0.05 mg/kg - Buprecare, Axience) before surgery. The surgical site was shaved, cleaned with antiseptic solution and a midline incision was made to expose the skull bone after local injection of Lidocaïne (4 mg/kg) (Laocaïne, Centravet). During all procedure, body temperature was maintained at 37°C by a thermostatically regulated heating pad, and eyes were covered with black tissue. The animal was placed in a stereotactic frame and holes were drilled at the injection sites. AAV suspensions were injected into the right hemisphere at two different locations in rats(2.6 mm ML, 6.8 mm AP and 3.1 mm ML, 7.2 mm AP from bregma) or at one location in mice (2.5 mm ML, 3.5 mm AP from bregma). In rats, for each injection site, 200 nl of viral vector (containing 0.2-8 × 10^15^ viral particles) was injected at three different depths (1100, 1350 and 1500 µm DV) with a micro-syringe pump controller (Micro4, World Precision Instruments) operating at a rate of 50 nl/min and a 10 µl Hamilton syringe. In mice, 1 µl of viral vector (containing 0.2-8 × 10^15^ viral particles) was injected at - 400 µm DV at a rate of 100 nL/min. At the end of surgery, animals were awakened with subcutaneous injection of Atipamazole (0.9 mg/kg) (Antidorm, Axience)

All animals were injected subcutaneously with Buprenorphine (0.05 mg/kg) one day after the intracranial injections (Buprecare, Axience). Electrophysiological recordings or craniotomy surgery for behavioral training in mice were performed at least one month after injections.

### *In vivo* extracellular recordings

Rats were anesthetized with a mixture of ketamine and medetomidine (40 mg/kg / 0.14 mg/kg). Pupils were dilated with tropicamide (Mydriaticum, Dispersa). A small craniotomy (5×5 mm square) was drilled above V1 in the right hemisphere. Before recording, tdTomato fluorescence was checked with a Micron IV retinal imaging microscope (Phoenix Research Laboratories). A 32-site µEcog electrode array (30 µm electrode diameter, 300 µm electrode spacing, FlexMEA36, MultiChannel Systems) was positioned over the transfected brain region for rats expressing MscL G22S or in a similar zone for control rats. After µEcog recordings, multi-electrode (MEA) recordings were performed with a 16-site silicon microprobe (electrode diameter 30 µm, spacing 50 µm, A1×16-5mm-50-703, NeuroNexus Technologies). The MEA probe was advanced 1100 µm into the cortex with a three-axis micromanipulator (Sutter Instruments, Novato, CA). The US transducer was coupled to the brain with a custom-made coupling cone filled with degassed water and US gel, and was positioned over the region of interest with a motorized stage. The probe and the US transducer were perpendicular for µEcog recordings and tilted at 45° for intracortical recording. The distance between the target in the cortex and the transducer was equal to the focal length of the transducer, as defined by custom-made US transducer coupling cones. Visual stimuli were generated by a white light-collimated LED (MNWHL4, Thorlabs) placed 15 cm away from the eye. Light irradiance at the level of the cornea was 4.5 mW/cm^2^. The µEcog and extracellular signals were digitized with a 32-channel amplifier and a 16-channel amplifier, respectively (model ME32/16-FAI-μPA, MultiChannel Systems). µEcog recordings were further analyzed with custom-developed Matlab scripts. MEA recordings were further analyzed with SpykingCircus software, and single-cell events were analyzed with custom-developed Matlab scripts. For µEcog recordings, response duration was calculated as the interval between the two time points at which the cortical evoked potential was equal to A/*e* (where A is peak depolarization and *e* is Euleur’s number). The peak depolarization of each channel was linearly interpolated to build pseudocolor activation maps. The activated area was defined as the area of the pseudocolor activation map over which peak depolarization exceeded the background noise level calculated as 2 times the standard deviation of the signal. The center of the response was estimated by weighting the peak depolarization of each electrode by its distance from other electrodes. For intracortical recordings, cell latency was estimated as the time between the stimulus onset and the maximum of the derivative of spike density function.

### Surgery for *in vivo* behavioral testing

Before surgery, mice were injected subcutaneously with Buprenorphine (0,05 mg/kg mouse) (Buprécare, Axience), and Dexamethasone (0,7 mg/kg) (Dexazone, Virbac). Animals were anesthetized with Isoflurane (5% induction, 2% maintenance, in air/oxygen mixture) and the head was shaved and cleaned with antiseptic solution. Next, animals were head-fixed on a stereotactic frame with an Isoflurane delivering system, eye ointment was applied and a black tissue was placed over the eyes. During all procedures when mice were under anesthesia, body temperature was maintained at 37°C by a thermostatically regulated heating pad. After a local injection of Lidocaïne (4 mg/kg) (Laocaïne, Centravet), an incision of the skin was made. Two screws were fixed in the skull, after a small craniotomy (approximately 5 mm x 5 mm) was drilled above V1 in the right hemisphere (0.5 mm steel drill) and cortex buffer was applied. The cortex was covered with a TPX plastic sheet (125 µm thick) and sealed with dental acrylic cement (Tetric Evoflow). For behavioral experiments, a metallic headbar (Phenosys) for head fixation was then glued to the skull on the left hemisphere with dental cement (FUJUCEM II). At the end of the surgery, animals were placed in a recovery chamber, with subcutaneous injection of physiological serum and ointment on the eyes (Ophtalon, Centravet). Buprenorphine was injected during post-surgery monitoring. Behavioral training in mice was performed at least 10 days after the surgical procedure.

### Mouse behavioral tests

C57BL6J mice were placed on a water restriction schedule, in which they received ∼ 0.5-1 mL of water per day until they reached approximately 80-85% of their weight with water supplied *ad libitum*. Mice were progressively habituated to drinking from a syringe, and head fixation and enclosure in a cylindrical body tube for the first five days. They were then trained to respond to a light stimulus by performing a voluntary detection task: licking a waterspout (blunt 18G needle, approximately 5 mm from mouth) in response to white light full-field stimulation (200 and 50 ms long) of the left eye (dilated with tropicamide, Mydriaticum Dispersa). Water (∼4 μL) was automatically dispensed 500 ms after the light was switched on, through a calibrated water system. The behavioral protocol and lick detection were controlled by a custom-made system, as previously described^43^. Visible light training lasted four days, with a typical training session lasting approximately 30 minutes and including 75-100 trials. After light stimulation training, four days (the first and second days were separated by a two-day break during the weekend) of US stimulation of the right hemisphere V1 were performed. During these four days, US stimulation was delivered for 50 ms at three different pressure values (0.2, 0.7 and 1.2 MPa). These pressure values were delivered each day but in a different order. Inter-trial intervals for light and US stimulation varied randomly and ranged between 10 and 30 s. The 15 MHz US transducer was coupled to the brain with a custom-made coupling cone filled with water and US gel, positioned over the region of interest with a motorized stage. We investigated the impact of light and US stimulation on mouse behavior, by assessing the success rate by counting the number of trials in which mice performed anticipatory licks (between the start of the stimulus and the opening of the water valve). The anticipatory lick rate was calculated by subtraction from the spontaneous lick rate (calculated in a 1 s time window before stimulus onset) and multiplication by the success rate. Lick latency was calculated by determining the time to the first anticipatory lick after stimulus onset.

### Immunohistochemistry and confocal imaging

Transduced retinas and brains were fixed by incubation in 4% paraformaldehyde (100496, Sigma-Aldrich) for 30 minutes for retinas, and overnight for brains. Brains were cryoprotected in 30% sucrose (84097, Sigma-Aldrich), and 50 µm thick sagittal slices were cut with a microtome (HM450, Microm). The slices displaying the highest levels of tdTomato fluorescence from each brain were selected for further immunohistochemistry and imaging. Retinas and sagittal brain cryosections were permeabilized by incubation in 0.5% Triton X-100 in PBS for 1 h at room temperature and then incubated in blocking buffer (PBS + 1% BSA + 0.1% Tween 20) for 1 h at room temperature. Samples were incubated overnight at 4 °C with a monoclonal anti-RBPMS antibody (1:500, Rabbit, ABN1362, Merck Millipore) for the retina, with a monoclonal anti-NeuN antibody (1:500; Mouse, MAB377, Merck Millipore) for brain sections in 0.5x blocking buffer supplemented with 0.5% Triton X-100. The sections were then incubated with secondary antibodies conjugated with Alexa Fluor (1:500; Molecular Probes) and DAPI (1:1000, D9542, Merck Millipore) for 1 h at room temperature. An Olympus FV1000 laser scanning confocal microscope with 20x objective (UPLSAPO 20XO, NA: 0.85) was used to acquire images of flat-mounted retinas and brain sections.

For the assessment of transduction efficiency, confocal images were processed with FIJI (ImageJ). RBPMS- and NeuN-positive cells were counted automatically with the *Analyze particles* FIJI plugin. MscL-tdTomato- and MscL-tdTomato-RBPMS/NeuN-positive cells were counted manually by two different users, with the CellCounter FIJI plugin. For the retina, quantification was performed by identifying the transfected area in each retina and acquiring confocal stacks in at least four randomly chosen regions of 0.4 mm^2^ per retina (Fig. E1). For V1 neurons, the sagittal brain slice with the largest *MscL*-expressing zone was selected for each animal. In some slices, tdTomato also diffused outside V1. A ROI in V1 was, therefore, manually defined and quantifications were performed in at least six randomly chosen regions of 0.4 mm^2^.

### US-induced tissue-heating simulations

When considering cell stimulation at higher US frequencies (15 MHz) than usually described in the US neuromodulation literature, it is essential to estimate thermal effects as they can become important. A three-fold process was used for this estimation: 1) simulation of the acoustic fields generated by the three transducers used, with realistic acoustic parameters, 2) verification that non-linear acoustics did not play an important role in heat transfer and 3) realistic simulations of the heat transfer and temperature rise induced at the focus by US in a linear regime for the parameters used in this study.

For non-linear simulations we used Matlab’s toolbox *kWave*, by defining the geometry of the transducer in 3D, and using the following parameters for the propagation medium (water): sound speed *c* = 1500 m s^-1^, volumetric mass *ρ* = 1000 kg m^-3^, non-linearity coefficient B/A = 5, attenuation coefficient *α* = 2.2 10^−3^ dB cm^-1^ MHz^-y^, and frequency power law of the attenuation coefficient y = 2 ^61^. We simulated quasi-monochromatic 3D wave-fields using long bursts of 50 cycles; this gave us both the maximum pressure field in 3D and the waveform at the focus. Simulations were calibrated by adjusting the input pressure (excitation of the simulated transducer) to reach the pressure at the focus measured in the water tank with the real transducers. The FWHM focal spot diameter in the *xy* plane was 4.36, 1.61 and 0.276 mm, and the major axis in the *xz* plane was 32.3, 20.6 and 3.75 mm long for the 0.5, 2.25 and 15 MHz transducers, respectively (Fig. 1b-d). Non-linear effects were evaluated by estimating the relative harmonic content of the waveform at the focus. In the 15 MHz focus transducer example in figure 1d, the experimental and simulated signals at the focal spot were compared and found to be highly concordant (Fig. E4a). Furthermore, the amplitude of the second harmonic is 19.8 dB below the fundamental (20.9 dB in the simulated case), meaning that if the fundamental energy is E, the second harmonic has energy E/95 (Fig. E4b). Therefore, we can reasonably neglect the non-linear effects in the calculations of the thermal effects, as they account for ∼1% of the energy involved. The same conclusions were drawn at 0.5 MHz and 15 MHz. The use of linear wave propagation approximations considerably decreased the computing cost of the simulations. Linear propagation simulations were conducted with the *Field II* toolbox in Matlab^62,63^, in monochromatic mode, with the same medium properties as *kWave* (water), to obtain the 3D maximum pressure fields. These maximum pressure fields were used to build a heating source term 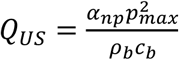, where *α*_*np*_ is the absorption coefficient of the brain at the considered frequency (59.04 Np m^-1^ at 15 MHz, calculated from *α*_*brain*_ = 0.21 dB cm^-1^ MHz^-y^ and y = 1.18), the brain volumetric mass *ρ*_*brain*_ = 1046 kg m^-3^, the brain sound speed *c*_*brain*_ *=* 1546 m s^-1 61,64^, and *p*_*max*_ is the 3D maximum pressure field. This source term was then used in the resolution of a Penne’s bioheat equation 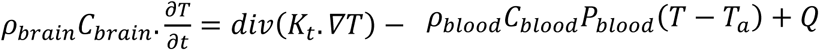 in *kWave*, where *C*_*brain*_ is the blood specific heat capacity (3630 J.kg^-1^ °C^-1^), *K*_*t*_ the brain thermal conductivity (0.51 W.m^-1^ °C^-1^), *P*_*blood*_ the blood density 1050 kg m^-3^, the blood specific heat capacity (3617 J.kg^-1^ °C^-1^), the blood perfusion coefficient (9.7 10^−3^ s^-1^), *T*_*a*_ the arterial temperature (37°C), and *Q = Q*_*US*_ + *ρ*_*brain*_ · *γ*_*brain*_ with *γ*_*brain*_ the heat generation of the brain tissue (11.37 W kg^-1^) ^64,65^. The initial condition for brain temperature was set to *T*_0_ = 37 °C.

This simulation corresponds to the worst case scenario regarding the temperature rise given: 1) that the acoustic propagation is simulated in water only, with a lower attenuation coefficient (2.2 10^−3^ dB cm MHz^-2^) than the brain (0.59 dB cm MHz^-1.27^), even if a part of the propagation occurs within the brain. *p*_*max*_ maps are, therefore, overestimated. 2) thermal absorption is simulated in brain tissue only, with a higher absorption coefficient (0.21 dB cm MHz^-1.18^) than water, even if a part of the maximum pressure field is actually located within the water of the acoustic coupling cone. *Q*_*US*_ is, therefore, slightly overestimated. We mapped the temperature in three spatial dimensions and time, and looked for the point of maximal temperature rise (Fig. E4 c-f).

### Statistical analysis

Statistical analysis was carried out with Prism software (Prism 7, GraphPad). All values are expressed and represented as means ± the standard error of the mean (SEM). The statistical tests performed are detailed in the figure legends. Data were analyzed in unpaired Welch’s *t*-tests (two-tailed).

## Data availability

The data supporting the findings of this study are available from the corresponding author upon reasonable request.

## Code availability

The custom Matlab codes are available from the corresponding author upon reasonable request.

## Acknowledgements

The authors would like to thank C. Joffrois, M. Valet, Q. Cesar, M. Desrosiers, S. Fouquet, P. Annic, M. Celik, Z. Raics for technical help and scientific advice. This work was supported by the European Research Council (ERC) Synergy Grant Scheme (holistic evaluation of light and multiwave applications to high-resolution imaging in ophthalmic translational research revisiting the Helmholtzian synergies, ERC Grant Agreement #610110), by the European Union’s Horizon 2020 research and innovation programme under grant agreement No. 785219 (Graphene Flagship Core 2) No. 881603 (Graphene flagship Core3), by the Foundation Fighting Blindness, *la Fédération des Aveugles de France*, Optic 2000, the city of Paris, *Région ile de France*, the *Agence Nationale de la Recherche* (ANR BrainOptoSight), and French state funds managed by the *Agence Nationale de la Recherche* (ANR) within *Programme Investissements d’Avenir, Laboratoire d’Excellence* (LABEX) LIFESENSES (ANR-10-LABX-0065) and *Institut Hospitalo-Universitaire* FOReSIGHT (ANR-18-IAHU-0001).

## Author Contributions

S.C, C.D. designed the experiments, S.C.,M.P., G.L.,I.A., J.L, R.G., E.B., J.D. carried out the experiments and analyzed the data, M.P., D.N., G.G., F.A., O.M., D.D., M.S., B.R. provided support for experiments, study design and data analysis, S.P., M.T., J.S. conceived the idea for this project and supervised the analysis of the data obtained. S.C., S.P., M.T., C.D, wrote the manuscript. All authors provided critical feedback on the research and the manuscript.

## Materials & Correspondence

Correspondence and requests for materials should be addressed to.

## Extended data figures

**Fig. E1.**
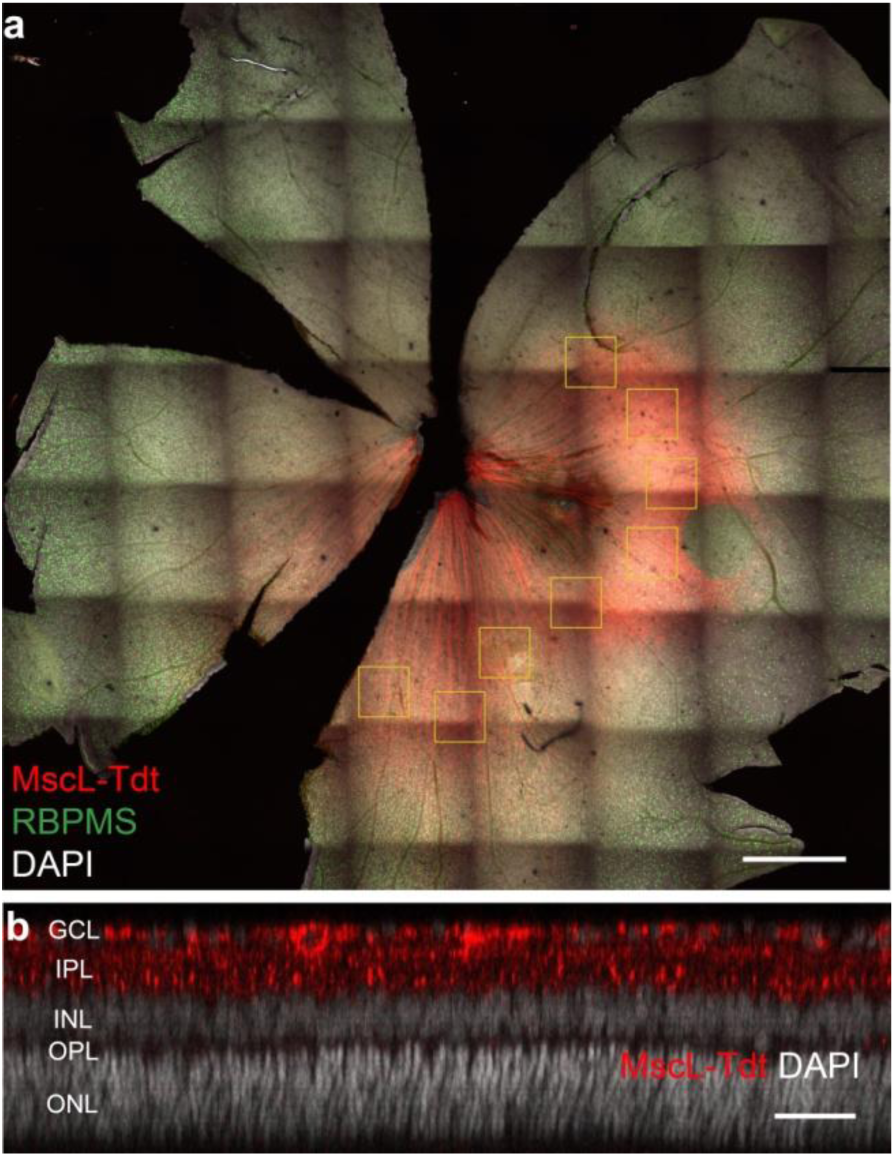
Retinal expression of MscL. (a) Whole-mount retina expressing MscL WT (red) and labeled with the RGC-specific anti-RBPMS antibody (green), with DAPI staining of the nucleus (white). Yellow boxes represent the 8 zones selected for the counting of MscL- and RBPMS-positive cells. (b) Optical section of a confocal stack showing MscL expression limited to the ganglion cell layer. The scale bars represent 1 mm in (a), 50 µm in (b).

**Fig. E2.**
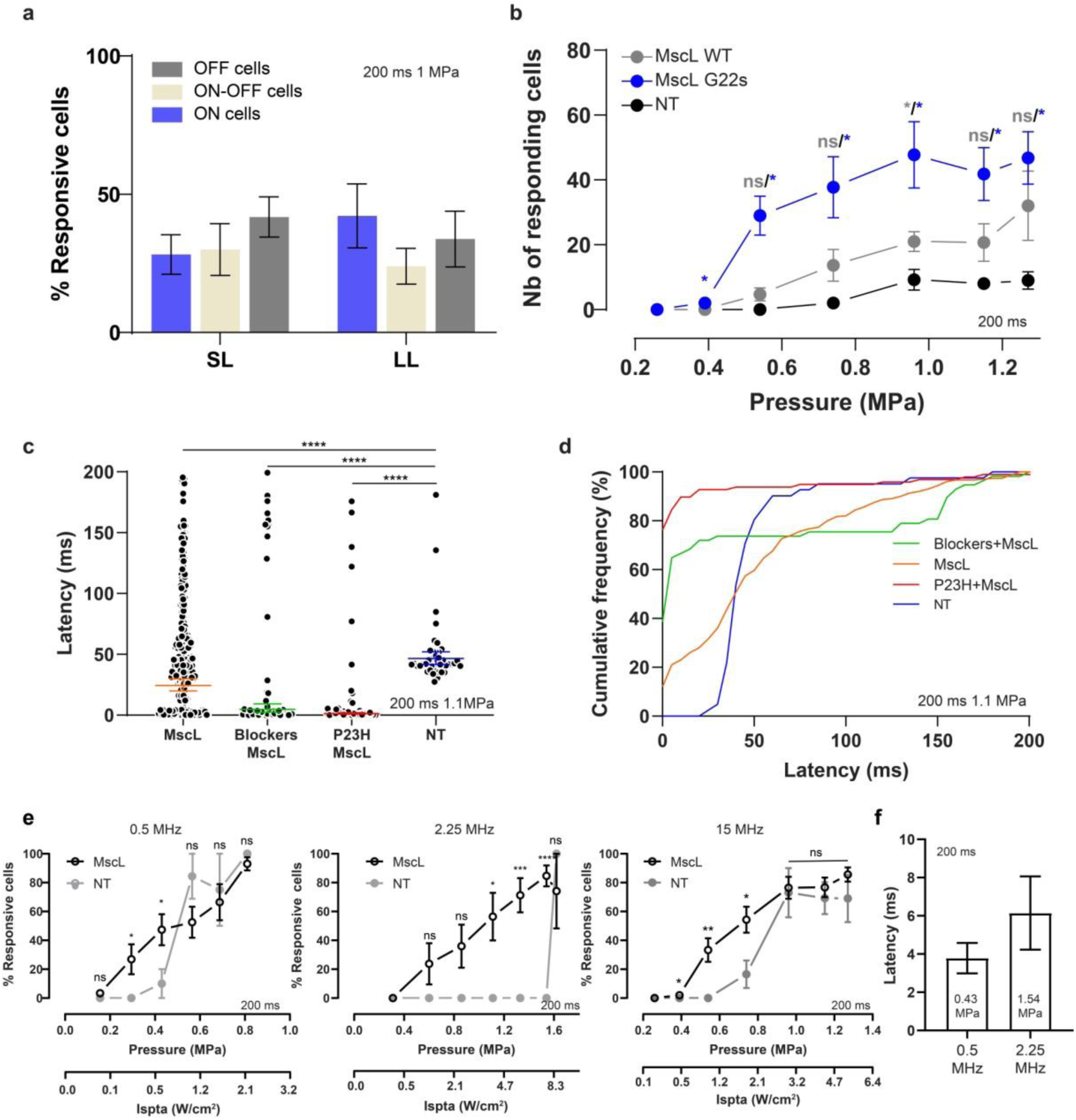
Retinal sonogenetic response characteristics for US stimuli of different frequencies. (a) Distribution of the different RGC cell types (ON, OFF, ON-OFF) among short (SL) and long latency (LL) responses in retinas (*n*=9) expressing MscL (WT and G22s form) following a 15 MHz US stimulus. (b) Numbers of RGCs responding to a 15 MHz stimulus of increasing acoustic pressure for MscL WT (*n*=3), MscL G22s (*n*=5) and NT (*n*=4) retinas (0.39 MPa: *, *p*=.0163; ns, 0.54 MPa: *p*=.1480*, *p*=.0168; 0.74 MPa: ns, *p*=.1334*, *p*=.312; 0.96 MPa: *, *p*=.0462*, *p*=.0279; 1.15 MPa: ns, *p*=.1617/*, *p*=.0145; 1.27 MPa: ns, *p*=.1580/*, *p*=.0144; unpaired *t* test between MscL WT and NT in gray and MscL-G22s and NT in blue). (c) Scatter plots and geometric means of RGC latencies in response to a 15 MHz US stimulus for MscL (*n*=300 cells), Blockers+MscL (*n*=57 cells), P23H+MscL (*n*=97 cells), and NT (*n*=41 cells) retinas (****, *p*<.0001, unpaired *t-*test on log-transformed values). (d) Cumulative frequency distribution of RGC latencies for MscL, Blockers+MscL, P23H+MscL, and NT retinas. (e) Percentage of cells responding to US stimuli (normalized against the maximum number of responsive cells in the experiment) of increasing acoustic pressure for 0.5 MHz (ns, *p*=.1661;*, *p*=.0292; *, *p*=.0260; ns, *p*=.8628; ns, *p*=.1316; ns, *p*=.7731; unpaired *t* test), 2.25 MHz (ns, *p*=.1474;ns, *p*=.0522; *, *p*=.0140; ***, *p*=.0005; ****, *p*<.0001; ns, *p*=.5000; unpaired *t* test) and 15 MHz (*, *p*=.0382;**, *p*=.0065; *, *p*=.0218; ns, *p*=.8628; ns, *p*=.5859; ns, *p*=.4223; unpaired *t* test) US. The lower *x* axis represents the corresponding acoustic intensity (Ispta). (f) Mean response latencies of SL cells for 0.5 and 2.25 MHz (*n*=9 and 8 retinas).

**Fig. E3.**
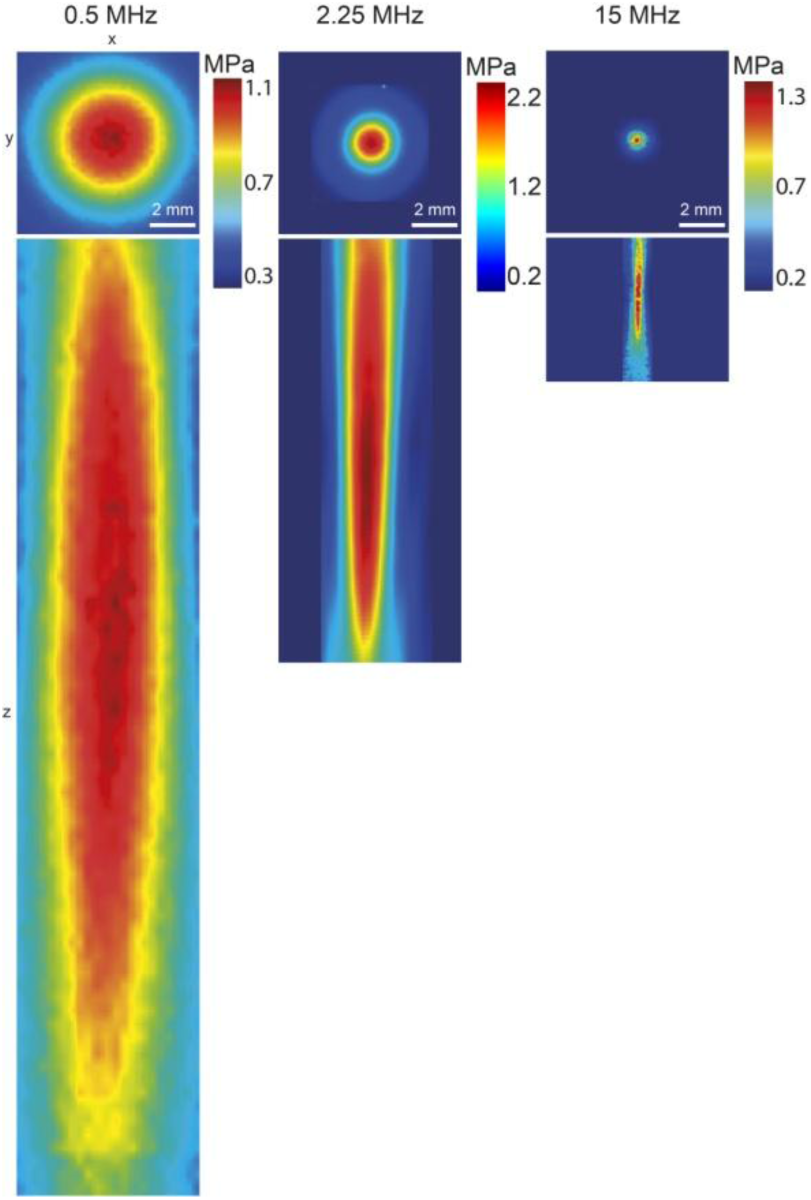
Experimentally measured US pressure fields near the focus for 0.5, 2.25 and 15 MHz focused transducers, measured in water. Color-coded pressure maps in the *xy* and *xz* planes, for 0.5, 2.25 and 15 MHz.

**Fig. E4.**
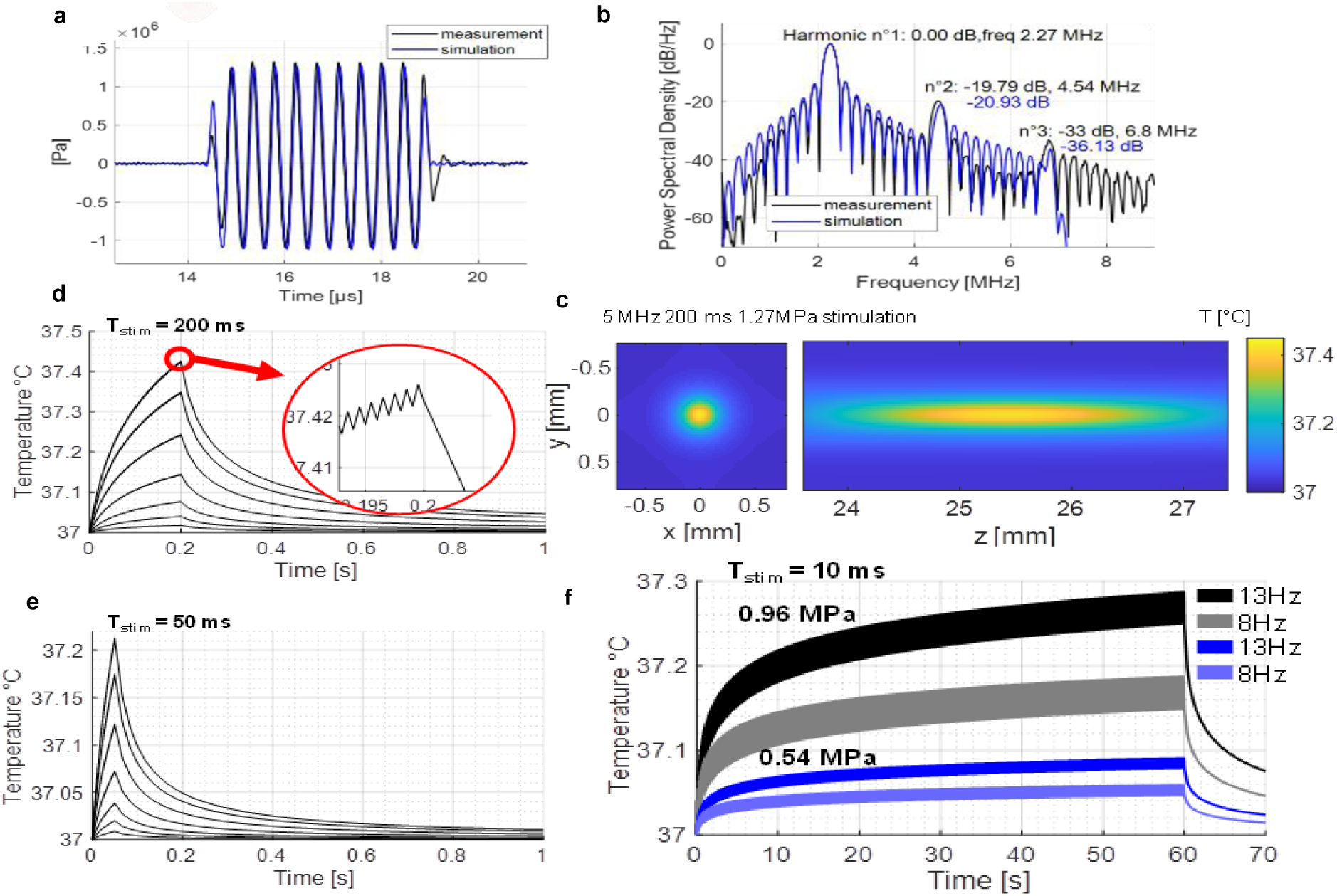
Simulated acoustic fields and temperature increases. (a) Comparison between a water tank measurement at the focus with a calibrated hydrophone (black) obtained with the 2.25 MHz transducer and reaching -1.11 MPa peak negative pressure, and a simulated waveform at the focus (blue) reaching the same negative pressure. The two waveforms match very well (0.42% error) ensuring a good match between our simulation setup and physical parameters. (b) Power spectral density of the measured (black) and simulated (blue) waveforms, showing that simulations can be used to estimate the importance of non-linear propagation. A second harmonic 20 dB below the fundamental indicates a factor of 100 in terms of energy, meaning that absorption can be calculated in a linear approximation. (c-f) Thermal simulations are performed in a two-fold process corresponding to a worst-case scenario (see methods): propagation in a water medium, and thermal absorption in a brain-mimicking medium. (h) 3D temperature map at the end of a 200 ms stimulation (at 15 MHz and 1.27 MPa). (d) Temperature rise at the focus for a 15 MHz 200 ms stimulation with the 7 pressures used in Fig. 1I (0.26, 0.39, 0.54, 0.74, 0.96, 1.15, 1.27 MPa). A zoom on the increasing curve reveals the fluctuations due to the 1 kHz on-off cycles. (e) Temperature rise at the focus for a 15 MHz 50 ms stimulation with the same 7 pressures. (f) Temperature rise at the focus for 15 MHz 10ms stimulations (1 kHz modulation) at a repetition rate of 8 Hz and 13 Hz (used in figure 3o), for focus pressures of 0.96 MPa and 0.54 MPa.

**Fig. E1.**
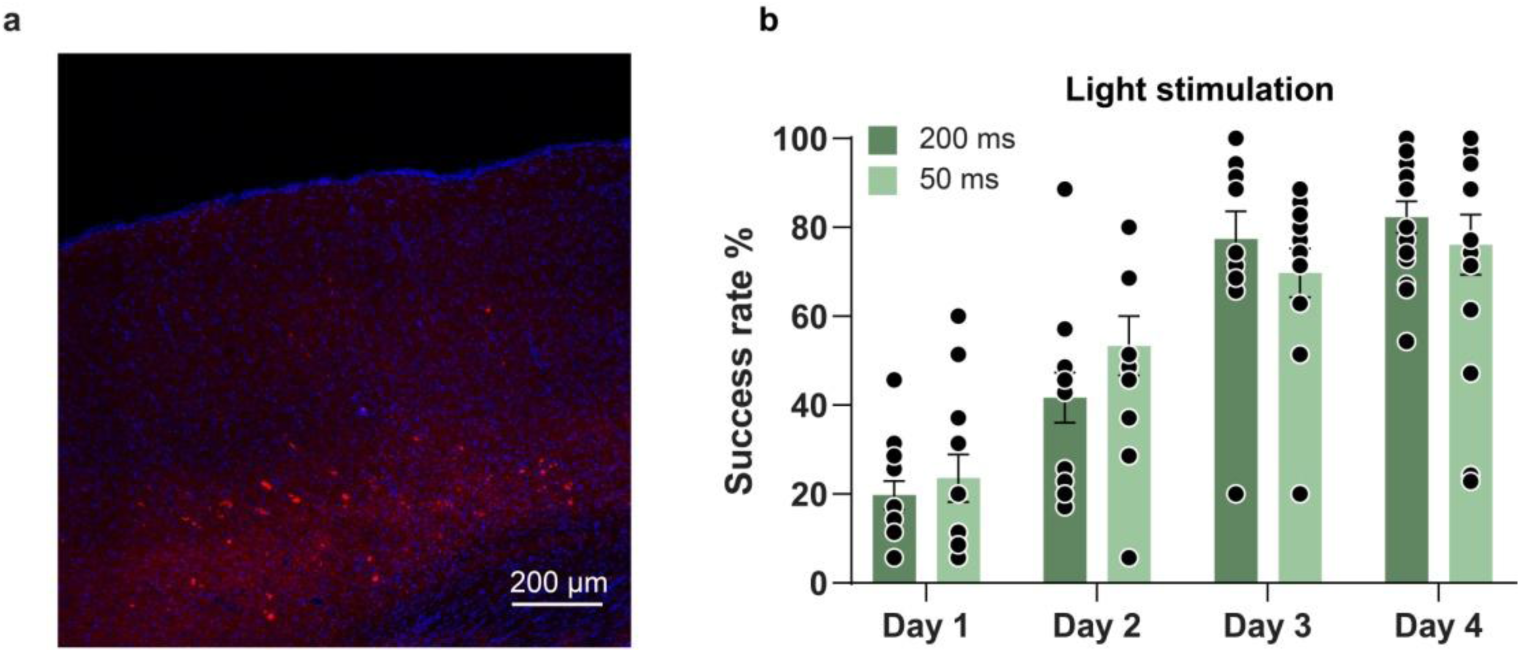
MscL G22S expression and light-associative training in mice. (a) Representative confocal stack projection of a sagittal brain slice expressing MscL G22s-tdTomato (red) and labeled with DAPI (blue). (b) Head-fixed and water-restricted mice were trained for four days to respond to a full-field stimulation of one eye (200 and 50 ms long) that preceding a water reward. Mice respond by licking before (anticipation — successful trial) or after the appearance of water. The success rate increased progressively and mice had learnt the task (upon 50 ms and 200 ms light stimulation) after four days of training.

